# A hormone-activated mobile RNAi pathway defends plant stem cells from virus infection

**DOI:** 10.1101/2022.12.18.520928

**Authors:** Marco Incarbone, Gabriele Bradamante, Florian Pruckner, Tobias Wegscheider, Wilfried Rozhon, Vu Nguyen, Ruben Gutzat, Thomas Lendl, Stuart MacFarlane, Michael Nodine, Ortrun Mittelsten Scheid

**Affiliations:** Gregor Mendel Institute of Molecular Plant Biology (GMI), Austrian Academy of Sciences, Vienna BioCenter (VBC), Vienna, Austria; Department of Agriculture, Ecotrophology, and Landscape Development, Anhalt University of Applied Sciences, Bernburg, Saxony-Anhalt, Germany; Research Institute of Molecular Pathology (IMP), Vienna BioCenter (VBC), Vienna, Austria; The James Hutton Institute, Invergowrie, Scotland, UK; Laboratory of Molecular Biology, Wageningen University and Research, 6700 AP Wageningen, The Netherlands

## Abstract

Stem cells are essential for the development and organ regeneration of multicellular organisms, so their infection by pathogenic viruses must be prevented. Accordingly, mammalian stem cells are highly resistant to viral infection due to dedicated antiviral pathways including RNA interference (RNAi) (*1, 2*). In plants, a small group of stem cells harbored within the shoot apical meristem (SAM) generates all postembryonic above-ground tissues, including the germline cells. Many viruses do not proliferate in these cells, yet the molecular bases of this exclusion remain only partially understood (*3, 4*). Here we show that a plant-encoded RNA-dependent RNA polymerase, after activation by the plant hormone salicylic acid, amplifies antiviral RNAi in infected tissues. This provides stem cells with RNA-based virus sequence information, which prevents virus proliferation. Furthermore, we find RNAi to be necessary for stem cell exclusion of several unrelated RNA viruses, despite their ability to efficiently suppress RNAi in the rest of the plant. This work elucidates a molecular pathway of great biological and economic relevance and lays the foundations for our future understanding of the unique systems underlying stem cell immunity.

## MAIN TEXT

Diseases caused by plant viruses are a constant threat to food and economic security worldwide, a reason for the extensive scientific investigation of plant-virus interactions. It remains poorly understood how viruses are excluded from stem cells in the SAM (*3*), even though this was first observed almost a century ago (*5*) and is common to many viral infections that efficiently spread throughout the rest of the plant. This particular antiviral capability of stem cells has been used to generate virus-free plants by tissue culture of meristems (*6*). After transition to flowering, SAM stem cells also generate floral organs containing the germline, so absence of virus in these cells is thought to play a key role in restricting vertical transmission of infection to the host progeny (*3*). Although the meristematic transcription factor WUSCHEL is involved in RNA virus exclusion from stem cells in *A. thaliana* (*4*) and RNAi and its suppression by viruses have also been implicated (*3, 7, 8*), the molecular mechanisms and dynamics of virus exclusion remain to be resolved.

To understand the events maintaining a virus-free niche in SAM stem cells, we challenged *A. thaliana* mutants lacking components of the RNAi pathway with Turnip mosaic virus expressing a fluorescent protein located at viral replication complexes (TuMV-6K2:Scarlet). Loss of RNA-Dependent RNA polymerase 1 (*RDR1*) caused TuMV to invade stem cells (**Fig. S1**). To document the dynamics of infection in wild type (WT) and *rdr1* we performed time-course experiments to assess virus propagation in the stem cell layers expressing a nuclear reporter expressed through the *pCLV3* promoter (**Fig. 1A**). This allowed a semi-quantitative approach and revealed temporary entry of TuMV in the top L1-L2 stem cell layers at 13-15 days post-inoculation (dpi), followed by subsequent exclusion (**Fig. 1B**). By contrast, *rdr1* mutants showed consistent virus infection of stem cells through time (**Fig. 1B**). This occurred even earlier in a double mutant with *rdr6* (**Fig. S2**), while in a *dcl2/dcl3/dcl4* (*dcl234*) mutant unable to generate small interfering (si)RNA we observed the highest levels of viral fluorescence in stem cells (**Fig. 1C**). These results, confirmed by *in situ* hybridization (**Fig. 1D**), portray a dynamic and layered RNAi antiviral network specifically protecting stem cells from infection. Moreover, RNAi did not exclude TuMV at the earliest time points, in accordance with observations with Cucumber mosaic virus (*4*). TuMV infection always caused loss of apical dominance, but while WT plants ultimately generated fertile flowers, *rdr1* mutants did not (**Fig. S3**), leading to sterility (**Fig. 1E**). RDR1 contributes to antiviral RNAi by increasing production of 21-22 nt-long virus-derived siRNA (vsiRNA) (*9*), presumably by generating double-stranded RNA (dsRNA) substrate for dicer enzymes. RDR1 significantly contributes to siRNA production from the whole TuMV genome (**Fig. 1F**; **S4A**,**C**,**D**) but, surprisingly, it does not affect overall TuMV accumulation (**Fig. 1F**; **S4B**). Finally, complementing *rdr1* with WT or RNA polymerization-deficient alleles of RDR1 provides evidence that dsRNA synthesis by this protein determines vsiRNA amplification (**Fig. 1G**), exclusion from stem cells (**Fig. 1H**) and fertility (**Fig. S3**).

**Figure 1:**
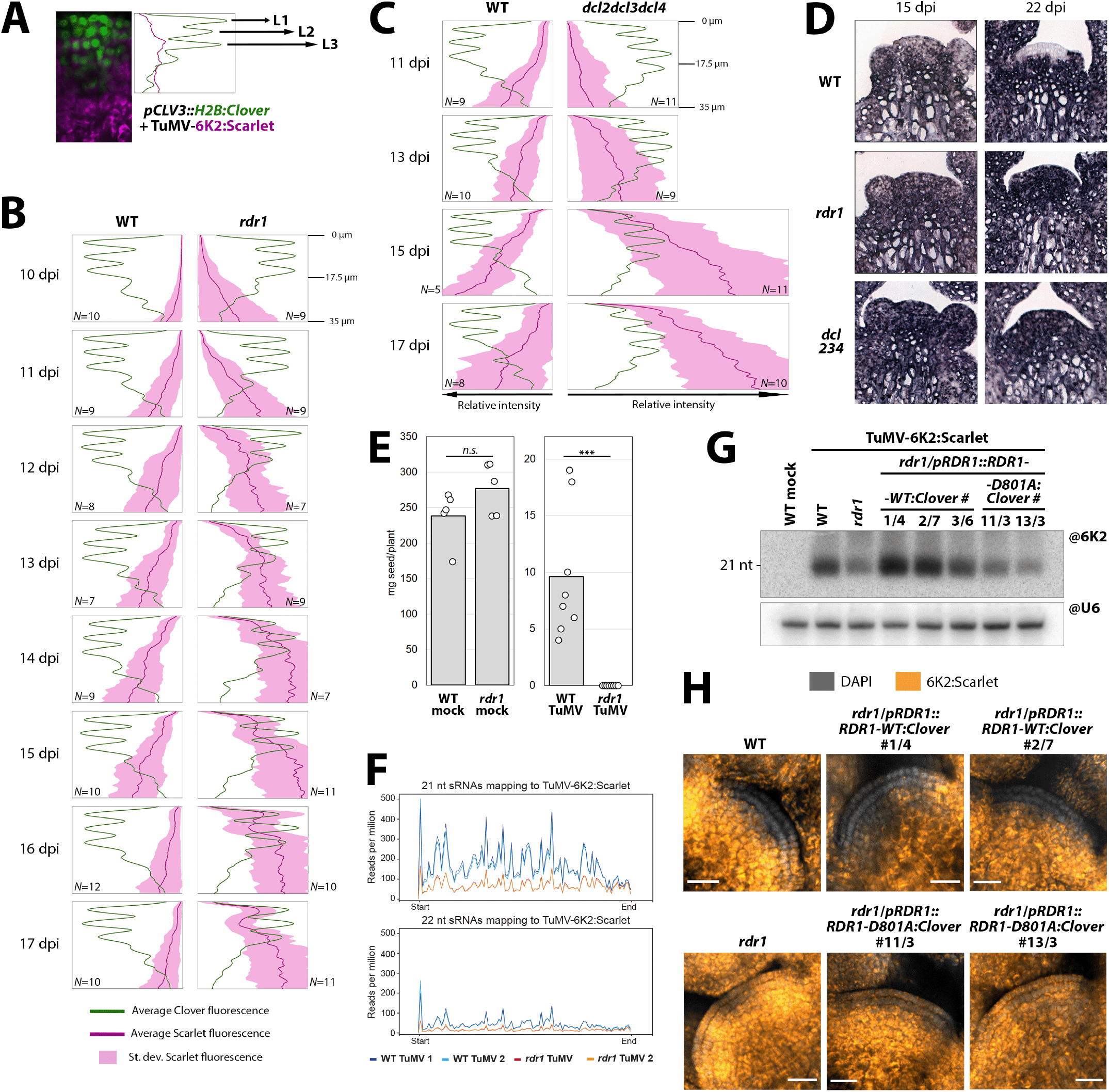
Arabidopsis RDR1 protects meristematic stem cells from TuMV infection through dsRNA synthesis and small RNA amplification. **(A)** Determination of virus entry into the stem cell area by quantification of fluorescence in plants expressing H2B:Clover (green) in SAM stem cell nuclei and infected with TuMV-6K2:Scarlet (magenta). **(B)** Fluorescence values as in (A) from the top 35 *μ*m of wild-type (WT) and *rdr1* SAMs between 10 and 17 days post-inoculation (dpi). Color legend at bottom, for simplicity Clover standard deviation not shown. *N*: number of meristems analyzed. **(C)** As in (B), fluorescence values in WT and *dcl234* triple mutants. **(D)** *In situ* hybridizations in vertical sections of WT, *rdr1* and *dcl234* meristems to detect TuMV RNA (purple) in infected plants at 15 and 22 dpi. **(E)** Seed production by mock and TuMV-infected WT and *rdr1* plants. Each data point represents progeny of one plant. *n*.*s*.: p>0.05; ***: *p*<0.001. **(F)** Distribution of vsiRNA along the TuMV-6K2:Scarlet genome, assessed by sRNA sequencing on duplicates of mock- and TuMV-infected apices (meristem and small flower buds). **(G)** Northern blot detection of TuMV-derived sRNA in *rdr1* expressing WT (*RDR1-WT:Clover*) or catalytically inactive (*RDR1-D801A:Clover*) alleles of *RDR1*. RNA was extracted from systemically infected leaves, snoRNA U6 is used as loading control. **(H)** Laser confocal microscopy of meristems from the lines in (G), 18 dpi. DAPI fluorescence in grayscale, Scarlet in orange-to-yellow, scale bar 20 *μ*m.

RNAi in plants has both local and remote, mobile silencing capabilities (*10*), the latter being well-documented for gene and transgene silencing but postulated indirectly for antiviral activity (*11, 12*). To assess whether RDR1 can act locally in stem cells, we generated *rdr1* lines expressing *pCLV3:RDR1*. These were able to restore TuMV exclusion (**Fig. 2A**). Interestingly, the exclusion zone was expanded to the whole *CLV3* promoter expression domain, establishing that RDR1 can prevent TuMV proliferation very efficiently and locally in stem cells. Yet, transcriptional reporters for the *RDR1* promoter showed that both in non- and TuMV-infected plants it drove expression in the lower meristem dome and the tissues below, but never in the core domain of stem cell virus exclusion (L1+L2 layers) (**Fig. 2B**). Along with reported expression in vasculature (*13*), this suggests that RDR1 is not produced in stem cells but prevents TuMV proliferation there through remote activity. Next, we asked whether RDR1 excludes TuMV from stem cells through *sensu stricto* antiviral RNAi or by regulation of gene expression through previously reported (*14*) and here confirmed host gene-derived siRNA (**Fig. S5**). To this end, we generated transgenic *rdr1* lines producing RDR1-independent antiviral siRNA (*siScar*) through a hairpin (**Fig. 2C, S6**). Production of *siScar* in stem cells of *rdr1* restored TuMV exclusion in a sequence-specific manner (**Fig. 2D**). Equally, production of *siScar* in subjacent non-stem cell tissues through the *RDR1* promoter yielded the same result (**Fig. 2E**), allowing to conclude that RDR1 excludes TuMV from stem cells by remotely providing viral RNA sequence information to the RNAi machinery, without the need for host gene-derived siRNA. As siRNAs are the mobile signal in RNAi (*12, 15*), we tested whether blocking 21-22 nt-long siRNA in cells expressing *RDR1* with the viral RNAi suppressor protein P19 (*15*) would stop the mobile signal and suppress the stem cell antiviral pathway. Surprisingly, this was not the case (**Fig. 2F**), in contrast to suppression of the pathway by P19 over-expression in all tissues - including stem cells (**Fig. 2G; Fig. S7**). This suggests that the mobile RDR1-dependent antiviral signal is either not 21-22 nt siRNA or a high load of 21-22 nt vsiRNA that requires a large amount of P19 to be blocked. Both possibilities explain why TuMV, which encodes a strong siRNA-sequestering RNAi suppressor (HC-Pro) (*16, 17*), cannot block this RNAi-based stem cell defense mechanism.

**Figure 2:**
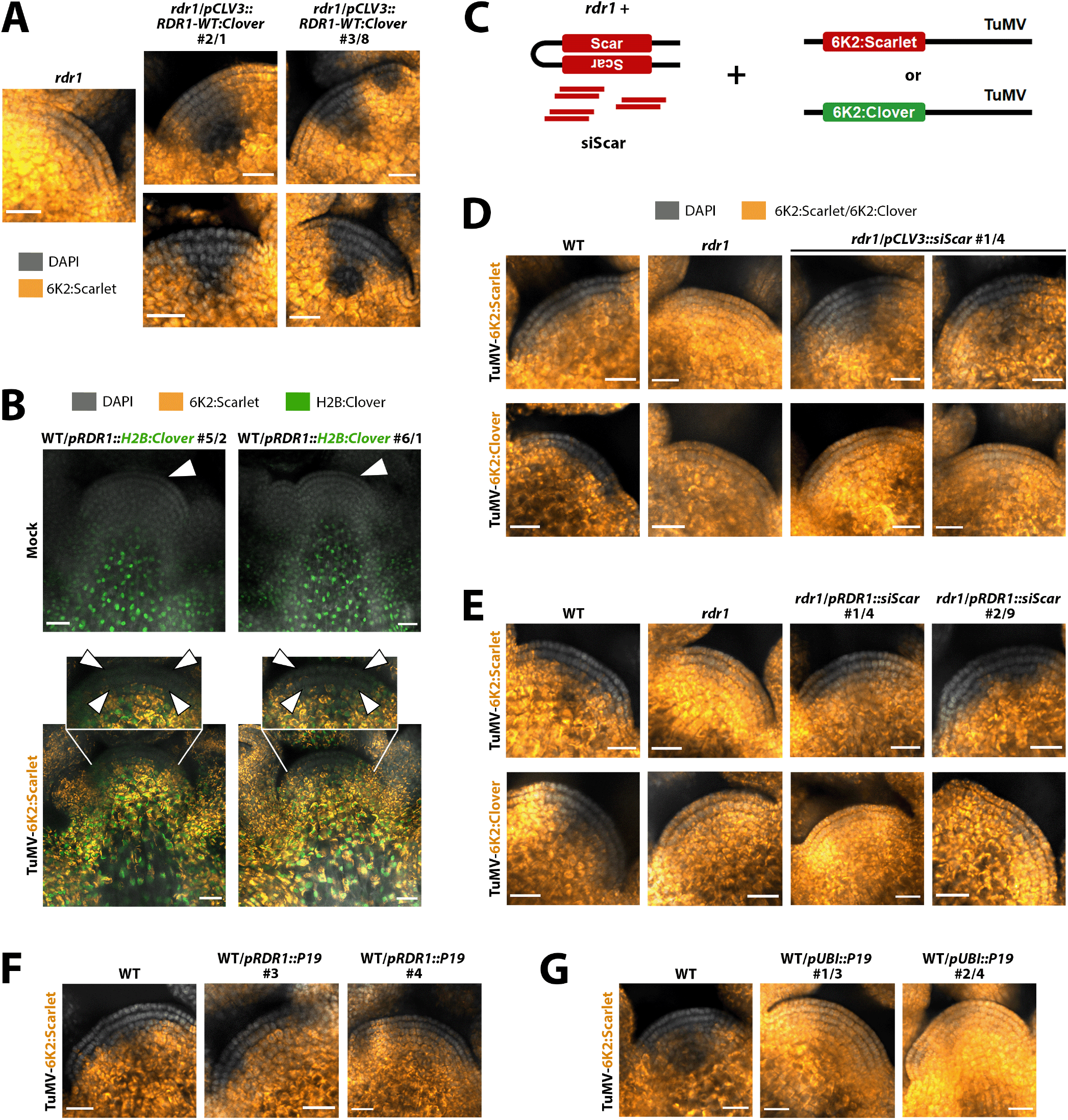
RDR1 immunizes the SAM stem cells at a distance by providing TuMV-specific small RNA. **(A)** Laser confocal microscopy of meristems from *rdr1* lines expressing *RDR1* through the stem cell-specific *pCLV3* promoter, infected with TuMV-6K2:Scarlet. **(B)** As in (A), of two independent transgenic lines expressing H2B:Clover (green) through the *pRDR1* promoter after mock or TuMV-6K2:Scarlet inoculation. Insets and white arrowheads: L1+L2 core virus exclusion zone. **(C)** Schematic representation of the *siScar* experiments: *rdr1* mutants generating Scarlet-specific siRNA (*siScar*) through a hairpin transgene are infected with TuMV containing the siRNA target sequence (TuMV-6K2:Scarlet) or not (TuMV-6K2:Clover). **(D)** Meristems of an *rdr1* mutant line expressing *siScar* in stem cells through the *pCLV3* promoter, infected with TuMV-6K2:Scarlet or TuMV-6K2:Clover. **(E)** As in (D), but lines expressing *siScar* through the *pRDR1* promoter. **(F)** Meristems of lines expressing P19 through the *pRDR1* promoter, after infection with TuMV-6K2:Scarlet. **(G)** As in (F), but of lines expressing P19 under the *pUBI* promoter. (A), (D), (E), (F), (G): DAPI fluorescence in grayscale, Scarlet or Clover in orange-to-yellow, scale bar 20 *μ*m.

*RDR1* expression is increased by salicylic acid (SA) in several crop species (*18*–*21*). SA is a key hormone in the activation of plant defenses against pathogens (*22*), including viruses (*23*), so we asked whether SA plays a role in TuMV exclusion from SAM stem cells. Indeed, TuMV completely invades stem cells of NahG plants (**Fig. 3A**) expressing a bacterial enzyme degrading SA (*24*). TuMV infection greatly increases SA accumulation in WT plants but not in NahG plants (**Fig. 3B**), and SA induction is required for *RDR1* upregulation upon infection (**Fig. 3C**). Increasing the steady-state amount of SA in plants lacking the SA-degrading *DMR6* gene (*25*) also leads to *RDR1* upregulation (**Fig. 3D, S8**). The TuMV-dependent SA response does not change in *rdr1* mutants (**Fig. 3D**), confirming that *RDR1* activation depends on SA and not vice-versa. As artificial overexpression of *RDR1* in NahG plants does not restore TuMV exclusion from stem cells (**Fig. S9**), transcriptional upregulation of RDR1 alone is not sufficient for SA-dependent virus exclusion. Therefore, either SA positively influences the RDR1 pathway by additional means such as protein activity/stabilization, or ubiquitous *RDR1* over-expression does not recapitulate SA-dependent induction. Our results do not exclude the possibility that SA acts through other molecular antiviral pathways. Nevertheless, *RDR1* is required for activation of SA to result in TuMV exclusion from stem cells, since *rdr1* mutants show an active SA pathway yet virus meristem invasion (**Figs. 1B, 3D**). Next, we asked whether SA activation is linked to stem cell exclusion of other virus species. We found that Turnip crinkle virus (TCV, family Tombusviridae) and Turnip yellow mosaic virus (TYMV, family Tymoviridae), species taxonomically distant from each other and TuMV (family Potyviridae), both elicit an SA response in WT Arabidopsis, albeit to different extents (**Fig. 3E**). TCV, the stronger inducer of SA, also upregulates *RDR1* expression (**Fig. S9**). *In situ* hybridizations revealed that both TYMV and TCV were excluded from SAM stem cells (**Fig. 3F,G**). Conversely, Tobacco rattle virus (TRV, family Virgaviridae), which infects meristems in *N. benthamiana* (*8*), did not elicit an SA response (**Fig. 3H**) and was not excluded from stem cells in *A. thaliana* (**Fig. 3I**). These results on four unrelated virus species therefore suggest that SA activation is correlated to the maintenance of virus-free SAM stem cells.

**Figure 3:**
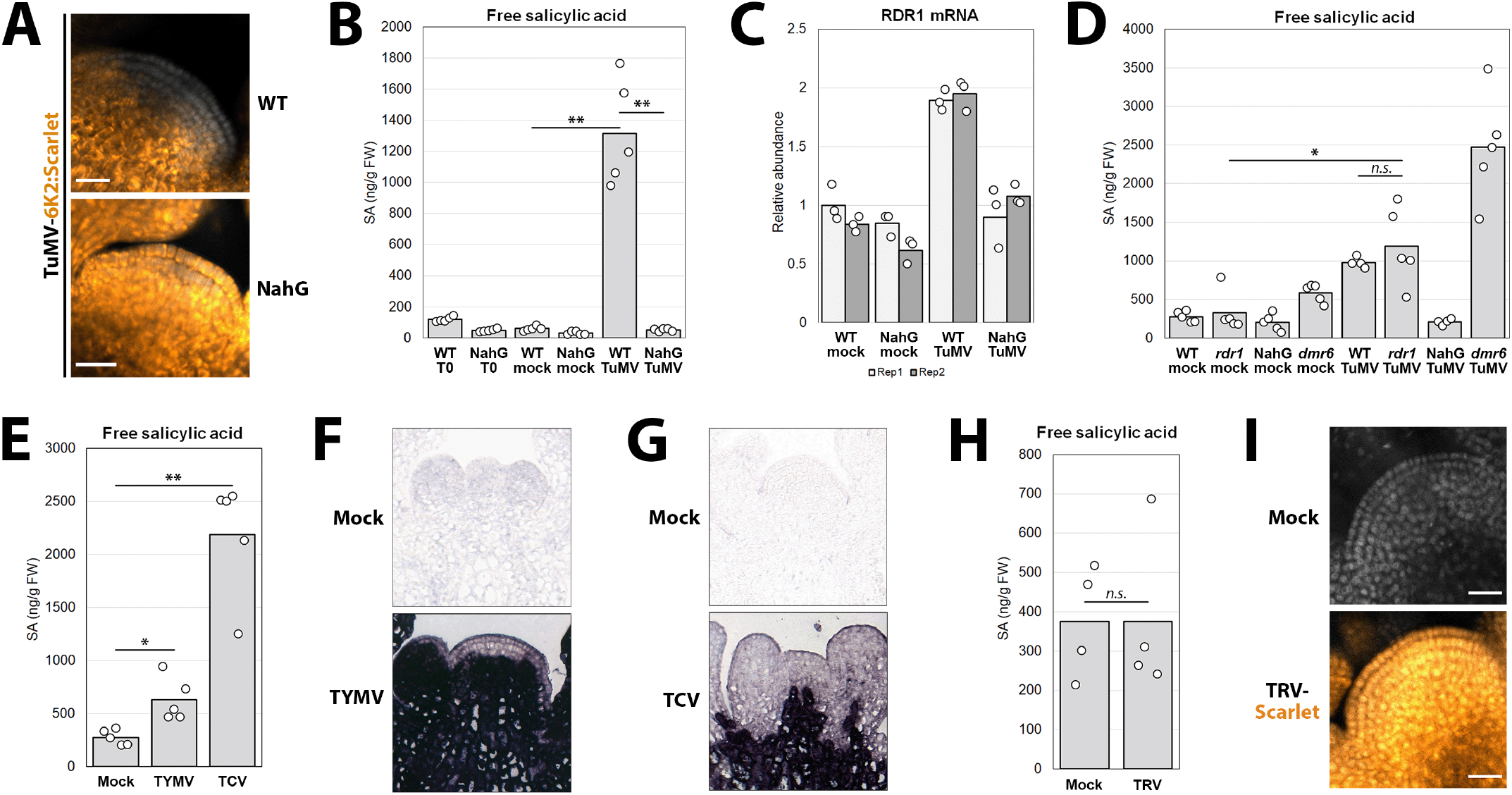
Increased salicylic acid (SA) production upon infection determines TuMV stem cell exclusion, increases *RDR1* expression and correlates with the exclusion of other virus species from stem cells. **(A)** Laser confocal microscopy of meristems from WT and SA-suppressing NahG plants infected with TuMV-6K2:Scarlet. **(B)** SA accumulation in WT and NahG plants before infection (T0) and after mock or TuMV-6K2:Scarlet inoculation. Each dot is a biological replicate: pool of tissues from 5-6 plants. **(C)** RT-qPCR on RNA from samples in (B) to assess *RDR1* mRNA accumulation. Each bar is a biological replicate, each dot is a technical replicate. **(D)** As in (B) on WT, NahG, *rdr1* and *dmr6* plants. **(E)** As in (B), but on WT plants infected with TYMV or TCV. Mock values are the same as in (D). **(F)** *In situ* hybridization to detect TYMV RNA (purple) in meristems of mock- or TYMV-inoculated WT plants, 15 dpi. **(G)** As in (F), to detect TCV RNA in meristems of mock- or TCV-inoculated WT plants, 15 dpi. **(H)** As in (B), but on WT plants infected with TRV-Scarlet. **(I)** As in (A), on WT plants after mock or TRV-Scarlet infection. (A), (I): DAPI fluorescence in grayscale, Scarlet in orange-to-yellow, scale bar 20 *μ*m. (B), (D), (E), (H): *n*.*s*.: *p*>0.05; *: *p*<0.05; **: *p*<0.01.

Next, we investigated whether RNAi and SA are necessary for stem cell exclusion of TCV and TYMV. TYMV and TCV can completely invade stem cells of *dcl234* mutants (**Fig. 4A,B**), indicating that small RNAs are required for exclusion. Furthermore, the expansion of the TCV exclusion zone over time is also dependent on small RNAs (**Fig. 4C**). Interestingly, neither *rdr1* nor NahG plants showed stem cell invasion, indicating that the SA/RDR1 pathway is not necessary for exclusion of these two viruses. TCV strongly induced SA/RDR1 production (**Fig. 3E, S9**), suggesting that this pathway is involved in – but not strictly necessary for – TCV exclusion from stem cells. These results suggest that for TCV and TYMV, either primary dicer products are sufficient and RDR1-dependent amplification of vsiRNA production is not required, or other RDR enzymes are involved and necessary here. Taken together, our observations establish that RNAi is essential in maintaining a virus-free SAM stem cell niche. This is remarkable, since as TuMV also TCV and TYMV encode for potent suppressors of RNAi (*26, 27*). Accordingly, *dcl234* mutants showed a modest increase in viral RNA accumulation, if any (**Fig. 4D**), indicating that host RNAi has little effect on virus replication/propagation in Arabidopsis plants at large. Strikingly however, in addition to ensuring stem cell exclusion, RNAi is required for TYMV- and TCV-infected plants to produce seeds (**Fig. 4E**). Whether virus stem cell exclusion and fertility are connected remains to be determined, but artificial exclusion of TuMV through *RDR1* or *siScar* expression in stem cells alone (**Fig. 2A, D**) does not rescue seed production in *rdr1* (**Fig. S10**), suggesting that virus exclusion *per se* is not sufficient to ensure fertility.

**Figure 4:**
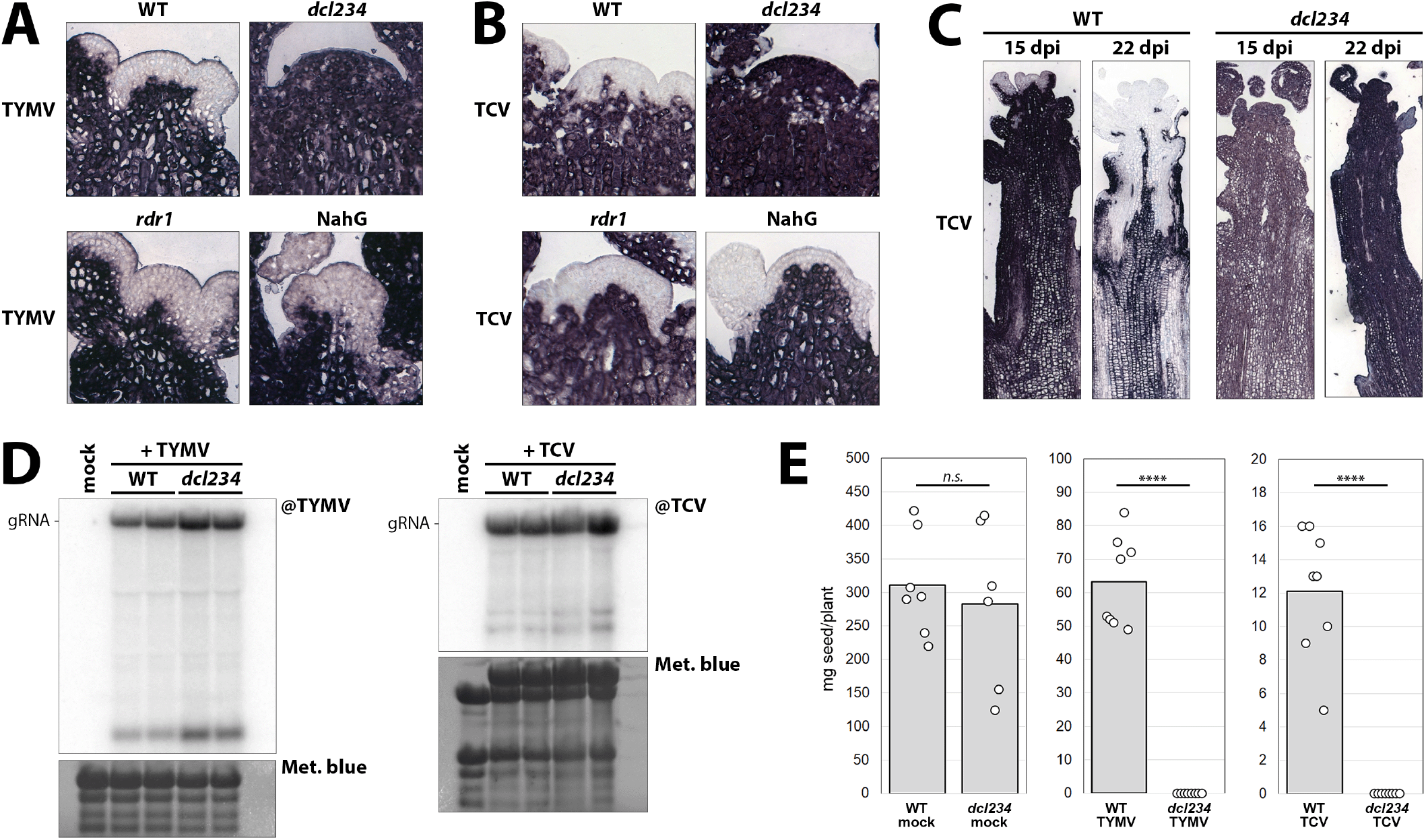
Small RNAS determine exclusion from stem cells of viral species unrelated to TuMV and are required for fertility of infected plants. **(A)** *In situ* hybridization to detect TYMV RNA in meristems of TYMV-inoculated WT, *dcl234, rdr1* and NahG plants, 15 dpi. Viral RNA results in blue-purple color. **(B)** As in (A) to detect TCV RNA in meristems of the same genotypes after TCV inoculation, 15 dpi. **(C)** As in (B), showing whole floral apices from WT and *dcl234* infected with TCV, 15 and 22 dpi. **(D)** Northern blot analysis of viral RNA accumulation in TYMV- and TCV-infected WT and *dcl234* plants, each in duplicate, from systemically infected leaves at 9 dpi. Methylene blue staining is used as loading control. **(E)** Seed production by mock-, TYMV- and TCV-infected WT and *dcl234* plants. Each data point is progeny of one plant. *n*.*s*.: *p*>0.05; ****: *p*<0.0001.

In summary, our study describes a broad-range antiviral RNAi pathway, which in the case of TuMV is non-cell autonomous and activated by salicylic acid, that maintains the vital plant SAM stem cells free of pathogenic viruses. Crucially, unlike RNAi in the rest of the plant, this pathway can successfully evade viral suppression, pointing to vital aspects of small RNA biology that remain to be elucidated. This work provides a robust molecular framework for a plant stem cell-specific defensive system of great biological and economic relevance.

## DATA AVAILABILITY

Additional and source data has been deposited on Zenodo at the following DOI: 10.5281/zenodo.7454454. This includes panels with complete confocal and *in situ* microscopy experiments, raw data from time course fluorescence quantifications, blotting, RT-qPCRs, SA measurements, complete sequences of virus and transgene plasmids.

## ACKNOWLEDGEMENTS

M.I. acknowledges funding from a Lise Meitner postdoctoral grant from the Austrian Science Fund (FWF M2921). O.M.S. is grateful for support by the Vienna Science and Technology Fund WWTF LS13-057, G.B. was supported by the Doctoral Program “Chromosome Dynamics” of the Austrian Science Fund (FWF W1238). M.I. and M.N. were supported by funding from the European Research Council under the European Union’s Horizon 2020 research and innovation program (Grant 637888 to M.N.). R.G. acknowledges support by the Austrian Science Fund (FWF I3687), S.M.F. is funded by the Scottish Government Rural and Environment Science and Analytical Services Strategic Research Programme. We would like to thank Pawel Pasierbek and Alberto Moreno Cencerrado at VBC BioOptics for assistance with microscopy, VBCF PlantS for assistance with plant work, VBC Histopathology and Magdalena Mosiolek for assistance with *in situ* hybridizations, VBCF NGS for small RNA library generation and sequencing, VBCF Protech for providing the pGGF-YFP seed selection plasmid, Eduardo Bejarano for the NahG and *sid2* seed stocks. We are grateful to lab members and GMI colleagues for ideas, feedback and discussions. Finally, we would like to thank the Vienna BioCenter in-house COVID-19 testing services for allowing us to safely carry out this research project during the coronavirus pandemic.

## AUTHOR CONTRIBUTIONS

M.I., G.B. and O.M.S. conceived the study and designed the experiments. M.I. cloned transgene and virus plasmids, generated and selected transgenic lines/mutants. M.I., G.B. and F.P. performed infections and confocal microscopy, G.B. and T.W. performed *in situ* hybridizations, T.L. and F.P. performed time-course fluorescence quantification and analysis. W.R. performed SA measurements and analyses. M.I. performed RNA extractions, RT-qPCRs and northern blotting, while V.N. and R.G. analyzed sRNA sequencing data. S.M.F. cloned the TRV1 plasmid. M.N. provided assistance and financial support in cloning and transgene generation. M.I. assembled the figures and wrote the manuscript, with the assistance of G.B. and O.M.S.

## MATERIALS AND METHODS

### Molecular cloning

All the binary plasmids used in this study were generated through Golden Gate assembly. Refer to Data Availability for the complete plasmid sequences. All transgenes for Arabidopsis transformation were assembled using the GreenGate system (*1*), by BsaI digestion (BsaI-HF-v2, New England Biolabs #R3733S) and T4 Ligase ligation (Thermo Scientific #EL0014) of entry vectors into binary plasmids. The entry vectors were made by amplifying the sequences of interest by PCR using Q5 HiFi DNA polymerase (NEB #M0491L) with primers containing the BsaI cut site in the appropriate orientation and the standard “sticky ends” corresponding to GreenGate A-F units (*1*), then ligating them into the pGGA-F GreenGate vectors or into the Golden Gate-ready pMiniT™2.0 (NEB #E1203S). Sequences containing BsaI cut sites (such as *AtRDR1*) were divided into several entry vectors, where final assembly would introduce silent mutations, preventing further digestion of the assembled products. Transgenes were assembled into the GreenGate pGreen-based pGGZ003 destination vector (*pCLV3::H2B:Clover*; *pUBI::H2B:Clover* and *pUBI::P19*) or pGGSun, a version of pSUN (*2*) we adapted for GreenGate cloning (*pCLV3::RDR1-WT:Clo*; *pCLV3::siScar*; *pRDR1::RDR1-WT:Clo*; *pRDR1::RDR1-D801A:Clo*; *pRDR1::siScar*; *pRDR1:P19*; *pUBI::RDR1-WT:Clo*; *pUBI::siScar*) (see **Table S1** for entry vectors used). The Golden Gate assembly-ready pSun, pGGSun, was obtained by amplifying (i) the pSun backbone adding BsaI sites and (ii) a ccdB selection cassette to insert between the BsaI sites. The two PCR products were then assembled with Gibson Assembly Mastermix (NEB #E2611). All RDR1 constructs contain the genomic sequence of RDR1 (AT1G14790). The catalytically inactive *RDR1-D801A* allele was generated by mutating aspartic acid 801 in the RDR1 protein to alanine. A corresponding mutation of the last aspartic acid in the conserved DxDxD triplet was shown to abrogate RNA polymerization capability in *A. thaliana* RDR2 and RDR6 (*3, 4*). The constructs using the *pRDR1* promoter do not only include the sequence upstream of *RDR1*, but also the sequence downstream of the *RDR1* gene, inserted downstream of the sequence of interest (*RDR1/H2B:Clover/siScar/P19*). The same is valid for constructs with the *pCLV3* promoter.

TuMV and TRV2 virus clones were generated by cloning segments of the viral genomes into pMini entry vectors, as described above, then seamlessly assembling them into pGGSun downstream of a 35S promoter and followed by a NosT terminator. In the case of TRV2-Scarlet (sequence of the PPK20 isolate), an HDV ribozyme was placed after the viral sequence to ensure cleavage for correct 3’ ending of the RNA. The mScarlet sequence, preceded by the PEBV CP subgenomic promoter, was inserted after the TRV CP-coding sequence. In the case of TuMV-6K2:Scarlet and TuMV-6K2:Clover, a sequence coding for the viral 6K2 protein (*5*) fused to Scarlet or Clover, respectively, and flanked by amino acid sequences cut by the viral proteases, was inserted into the polycistronic TuMV sequence (UK1 isolate) between the P1- and HC-Pro-coding sequences. The TRV1 plasmid (p1586 - pCB-TRV1) was generated by cloning the cDNA from TRV1 isolate PPK20 into binary vector pDIVA (*6*), between the 35S promoter and the HDV ribozyme, by blunt ligation into the PCR-amplified backbone.

### Plant material

For ease of interpretation, in this manuscript wild-type (WT) is used to refer to *Arabidopsis thaliana* Col-0 ecotype plants, which is also the genetic background of all mutants. Arabidopsis mutant lines *rdr1-1* (*7*), *rdr6-15, rdr1-1/rdr6-15* (*8*), *dcl2-1/dcl3-1/dcl4-2* (*9*), NahG (*10*) and *dmr6-2* (*11*) were previously described (See **Table S2** for stock and genotyping information). Genotyping was performed by standard PCR of leaf DNA extracts. Transgenic Arabidopsis lines were generated by transforming *A. tumefaciens* GV3101 with the plasmid of interest and using the resulting cultures to perform floral dip. The transformants were selected in the appropriate manner (antibiotic resistance or seed coat fluorescence) and propagated to the third generation after transformation, when seed stocks homozygous for the transgene were selected and further used for infection experiments. The lines used for time-course experiments (*pCLV3::H2B:Clover* in Col-0, *rdr1, rdr6, rdr1/rdr6* and *dcl2/dcl3/dcl4* backgrounds) were obtained by crossing a Col-0/*pCLV3::H2B:Clover* line with *rdr1-1/rdr6-15* or *dcl2-1/dcl3-1/dcl4-2* and selecting the various mutant combinations by genotyping. All other transgenics were obtained by directly transforming the genotypes in question. In all infection experiments, plants were grown on soil at 12 h/12 h day/night cycles until infection, when they were moved to 16 h/8 h long day conditions to induce flowering. Plants were infected 3.5/4 weeks after germination (TuMV, TCV, TYMV) or 2 weeks after germination (TRV).

### Virus infection, tissue sampling and meristem preparation

Inoculum of TuMV-6K2:Scarlet and TuMV-6K2:Clover was obtained by inoculating *N. benthamiana* plants with *A. tumefaciens* cultures containing the respective plasmids as previously described (*12*) followed by harvesting and freezing the systemically infected leaves. Inoculum of TCV and TYMV was obtained by harvesting and freezing Arabidopsis leaves systemically infected after rub inoculation. Inoculum of TRV-Scarlet was obtained by harvesting and freezing Arabidopsis leaves systemically infected after inoculation of *A. tumefaciens* cultures containing TRV1 and TRV2-Scarlet plasmids as previously described (*12*). During infection experiments, inoculum was prepared by grinding frozen plant tissue in liquid nitrogen with mortar and pestle, then resuspending the powder in 50 mM sodium phosphate buffer pH 7.2, 0.2% sodium sulfite. After incubating on a wheel at 4°C for 10 min, the homogenate was centrifuged at 1000 g for 2 min, the supernatant kept on ice and used as inoculum. Plants were sprinkled with Celite 545 (Merck), a cotton swab was dipped in the inoculum and used to gently rub the leaves, 5-6 leaves per plant. For molecular analysis, tissues were harvested at 8-9 dpi (systemic leaves) or 15-16 dpi (inflorescence apices), frozen and stored at -70°C. Each sample is a pool of tissues from 4-5 plants. For meristem preparations, the main inflorescence of each plant was removed and dissected under a light microscope until only the smallest flower buds and shoot apical meristem remained along with 1-2 mm of stem. Unless indicated otherwise in figures or figure legends, meristems were generally sampled at 15-18 dpi, depending on the experiment, with the exception of the *pRDR1::H2B:Clover* experiments at 12-13 dpi. Precise time points are indicated in the additional microscopy data (see Data Availability). If meristems were to be observed by confocal microscopy, the dissected meristems were incubated 40 min in fixing solution (*13*) (1x MTSB, 2% paraformaldehyde, 0.1% Triton-X) at 37°C, then stored in MTSB at 4°C for a maximum of 10 days. 3-4 days before observation, the meristems were incubated in ClearSee (10% w/v xylitol, 15% w/v sodium deoxycholate, 25% w/v urea) at 4°C, with the addition of 10 mg/L DAPI the day before observation. If the meristems were to be used for *in situ* hybridizations, they were incubated after dissection over-night at 4°C in fixing solution (4% formaldehyde, 50% ethanol, 5% glacial acetic acid, 1x PBS) and dehydrated by changing the buffer to 50% then 70% ethanol in 1x PBS.

### Confocal microscopy and image analysis

Meristems were mounted on glass slides in ClearSee and imaged with a Zeiss LSM880 laser confocal microscope. The following laser wavelengths were used: 405 nm for DAPI, 488 nm for Clover, 561 nm for Scarlet. Further image processing was carried out with FIJI/ImageJ. For single meristem image assembly, images were cropped, rotated if necessary, split into single channels, LUTs were assigned (grayscale for DAPI, green for Clover, OrangeHot for Scarlet or Clover in **Fig. 2D,E**) and brightness/contrast were adjusted. For Scarlet fluorescence signal, brightness was regulated, until the high-signal zones were in yellow color, with the same settings for all genotypes. The single channel images were then merged and a 20 *μ*m scale bar was added (**Fig. S1**). For time course quantification experiments (**Fig. 1B,C**; **Fig S2**), 7-12 meristems were imaged per genotype/time-point without changing laser intensities within an experiment. Images were then analyzed with FIJI (*14*) using a macro developed for this task (**Supplementary File 1**): with meristems oriented vertically, an equally wide vertical section of each was selected for Clover and Scarlet fluorescence quantification, one measurement every 149 nm. The data were then imported into Microsoft Excel spreadsheets. Since differences in sample depth and degree of clearing caused differences in absolute fluorescence between meristems, the Clover fluorescence values in each meristem were converted to values on a 0-100 scale. The corresponding Scarlet values were normalized for each data point. These normalized values were then used to calculate the plotted average and standard deviation.

### *In situ* hybridization

Meristems, after being prepared as described above, were stained in 1% w/v eosin in 70% ethanol then infiltrated with xylene substitute and paraffin in a Diapath Donatello I tissue processor, after which they were cast into paraffin blocks using a Sakura Tissue Tek TEC5 (approximately 20 meristems/block). The blocks were then cut into 2 *μ*m-thick sections that were transferred onto glass microscopy slides, which were screened for ones containing central sections of meristems. DIG-labelled RNA probes were generated with DIG RNA Labeling Kit T7/SP6 (Roche #11175025910), see **Table S3** for the primers used to generate the DNA templates. *In situ* hybridization was then performed as previously described (*15, 16*), with minor variations, all solutions being prepared with DEPC-treated water. Slides were twice incubated 10 min in Histo-Clear II (National Diagnostics #HS-202), 5 min in 100% ethanol twice and rehydrated through serial passages in 90%, 70%, 50% and 30% ethanol, then in Tris-EDTA pH 7.5. Sections were then treated with Proteinase K (Roche #3115836001), washed in 1x PBS, incubated 10 min in 4% paraformaldehyde, dehydrated through serial ethanol washes and air-dried. After probe denaturation for 3 min at 80°C, hybridization with 50-100 ng DIG-labelled probes per slide was carried out O/N at 50°C in 150 *μ*l hybridization solution: 50% formamide, 10 mM Tris base, 300 mM NaCl, 5 mM EDTA, 10 mM Na_2_HPO_4_, 1x Denhardt’s solution (Sigma Aldrich #D2532-5ML), 10% dextran sulphate, 0.5 *μ*g/*μ*l tRNA (Roche #10109517001). Slides were briefly washed in 2x SSC, then incubated in 0.2x SSC for 2 h at 55°C and treated with RNase A (Thermo Scientific #EN0531) at 37°C for 30 min, then 1 h in 0.2x SSC at 55°C. Slides were washed 10 min in washing buffer and incubated 1 h in blocking buffer (both Roche #11585762001). Anti-DIG antibody was added (Roche #11093274910 - 1:1500 dilution) and incubated for 1 h 45 min at room temperature, washed for 1 h, incubated in TNM5 (100 mM Tris pH 9.5, 100 mM NaCl, 5 mM MgCl_2_) three times for 2 min, then O/N in TNM5 with 10% w/v polyvinyl alcohol, 10 *μ*l/ml NBT/BCIP (Roche #11697471001). Slides were mounted with Aqua-Poly/Mount (Polysciences #18606-20) and scanned with a Pannoramic 250 slide scanner at 40 x magnification.

### Salicylic acid quantification

For SA quantification, 4-5 replicates of each genotype/virus were collected, each replicate being a pool of systemically infected tissue from five plants. Tissues were frozen, pulverized and stored at -70°C. Aliquots of tissue were weighed, ground with glass beads and 1 ml of 80% acetonitrile (Sigma-Aldrich #34881) and 50 *μ*l internal standard (5-fluorosalicylic acid, 1 mg/l) were added per sample. The resulting solution was then vortexed and placed in a shaker at room temperature for 1 h at 1400 rpm shaking speed. Samples were centrifuged 5 min at 13000 rpm and supernatant was transferred to new tubes. After drying most of the liquid with a vacuum pump, quantification was performed by HPLC as previously described (*17*), with the difference that a Nucleodur 100-5 NH2 125x4 mm column (Macherey-Nagel, #760730.40) and an eluent consisting of 8.5% acetonitrile and 25mM formic acid pH 4 were used (*18*). Data was analyzed and plotted with Microsoft Excel, significance was assessed through standard pairwise t-student tests, two-tailed, assuming unequal variance.

### Northern blotting and RT-qPCR

RNA extraction was performed with TRI Reagent (Zymo Research #R2050-1-200). Briefly, flash-frozen plant tissues were pulverized with glass beads, 1 ml TRI Reagent and after clearing 300*μ*l chloroform were added. After shaking and centrifugation, one volume isopropanol was added to the aqueous phase and incubated at least 1 h on ice. After centrifugation, the pellet was washed with 80% ethanol, dried, resuspended in RNase-free water and the RNA concentration measured. RNA was stored at -20°C. Small RNA northern blotting was performed as previously described (*19*) on 10-50 *μ*g RNA, using standard BioRad PAGE system for electrophoresis and EDC chemical cross-linking (Sigma Aldrich #E7750) onto Hybond NX nylon (GE Healthcare #RPN203T). Membranes were probed with *α*-^32^P-CTP-labelled (Agilent #300385) PCR products (@6K2, @Scarlet) or *γ*-^32^P-ATP-labelled (Thermo Scientific #EK0031) DNA oligonucleotides (@U6), hybridizing overnight at 42°C in 1 mM EDTA, 7% SDS, 500 mM sodium phosphate pH 7.2. After three washes of 15 min at 45°C in 2% SDS, 2x SSC, membranes were exposed to phosphor screen and signals revealed by an Amersham Typhoon. For high molecular weight northern blotting to detect viral RNA, 5 *μ*g RNA was initially denatured by incubating with 15% v/v deionized glyoxal at 50°C for 1 h. Samples were then run in a 1% agarose gel in 20 mM sodium phosphate pH 7.2, capillary transfer to nylon membrane was performed overnight followed by UV cross-linking. After staining with methylene blue, membranes were probed with *γ*-^32^P-ATP-labelled DNA oligonucleotides (@TCV, @TYMV) as described above. For RT-qPCR quantification, 5 *μ*g RNA was treated with TURBO™ DNase (Invitrogen #AM2238) and 500 ng of this was used for cDNA synthesis with oligo-dT primer using RevertAid H Minus First Strand cDNA Synthesis (Thermo Scientific #K1632). qPCR on cDNA was performed with FastStart Essential DNA Green Master kit (Roche #06402712001) using a Roche LightCycler 96 and corresponding proprietary software. Expression levels of *RDR1* and *PR1* were normalized to housekeeping gene *AtSAND* (AT2G28390), while levels of TuMV were normalized to *AtGAPDH* (AT1G13440). Data were analyzed and plotted with Microsoft Excel. See **Table S3** for primer sequences.

### Small RNA sequencing and analysis

sRNA libraries were generated with QIAgen miRNA library kit (QIAgen #331502) according to manufacturer’s instructions. Following quality control they were sequenced on an Illumina HiSeq2500 with HiSeq V4 reagents, single-read 50 read-mode, all steps performed by the VBCF Next Generation Sequencing Facility. Prior to analyzing the sequencing data, adapters were removed from sRNA library data by using cutadapt v1.18, selecting read length from 18 to 26 nt. Processed reads were aligned to the Arabidopsis genome (TAIR10) and TuMV-Scarlet sequence using bowtie2 v2.3.5 (*20*), (i) allowing unique mapping to the TuMV-6K2:Scarlet sequence to assess the proportion of viral sRNAs and (ii) allowing 1000 times multi-mapping for gene-derived sRNA enrichment analysis. The aligned reads from multi-mapping were sorted by size (21 nt, 22 nt and 24 nt) for further analysis. The small RNA metaplots were generated by using Deeptools v.3.3.1 (*21*) with “bamCoverage” adding “CPM” parameter. The annotation of small RNAs to genes was done by using featureCounts (*22*) with Araport 11 annotation. DESeq2 (*23*) was used to analyze small RNA enrichment on genes with a cutoff of p.adj. < 0.05, log2 fold change > |1| and > 10 reads in both replicates. Visualization of the data was done by using the packages tidyverse (*24*) and ggplot2 (*25*).

## REFERENCES

1. X. Wu, V. L. Dao Thi, Y. Huang, E. Billerbeck, D. Saha, H.-H. Hoffmann, Y. Wang, L. A. V. Silva, S. Sarbanes, T. Sun, L. Andrus, Y. Yu, C. Quirk, M. Li, M. R. MacDonald, W. M. Schneider, X. An, B. R. Rosenberg, C. M. Rice, Intrinsic Immunity Shapes Viral Resistance of Stem Cells. Cell. 172, 423-438.e25 (2018).

2. E. Z. Poirier, M. D. Buck, P. Chakravarty, J. Carvalho, B. Frederico, A. Cardoso, L. Healy, R. Ulferts, R. Beale, C. Reis e Sousa, An isoform of Dicer protects mammalian stem cells against multiple RNA viruses. Science 373, 231–236 (2021).

3. G. Bradamante, O. Mittelsten Scheid, M. Incarbone, Under siege: virus control in plant meristems and progeny. Plant Cell. 33, 2523–2537 (2021).

4. H. Wu, X. Qu, Z. Dong, L. Luo, C. Shao, J. Forner, J. U. Lohmann, M. Su, M. Xu, X. Liu, L. Zhu, J. Zeng, S. Liu, Z. Tian, Z. Zhao, WUSCHEL triggers innate antiviral immunity in plant stem cells. Science 370, 227–231 (2020).

5. C. W. Bennett, The relation of viruses to plant tissues. Bot. Rev. 6, 427–473 (1940).

6. A. Panattoni, A. Luvisi, E. Triolo, Elimination of viruses in plants: twenty years of progress. Spanish J. Agric. Res. 11, 173 (2013).

7. F. Schwach, F. E. Vaistij, L. Jones, D. C. Baulcombe, An RNA-Dependent RNA Polymerase Prevents Meristem Invasion by Potato Virus X and Is Required for the Activity But Not the Production of a Systemic Silencing Signal. Plant Physiol. 138, 1842–1852 (2005).

8. A. M. Martin-Hernandez, D. C. Baulcombe, Tobacco Rattle Virus 16-Kilodalton Protein Encodes a Suppressor of RNA Silencing That Allows Transient Viral Entry in Meristems. J. Virol. 82, 4064–4071 (2008).

9. X.-B. Wang, J. Jovel, P. Udomporn, Y. Wang, Q. Wu, W.-X. Li, V. Gasciolli, H. Vaucheret, S.-W. Ding, The 21-nucleotide, but not 22-nucleotide, viral secondary small interfering RNAs direct potent antiviral defense by two cooperative argonautes in Arabidopsis thaliana. Plant Cell. 23, 1625–38 (2011).

10. X. Chen, O. Rechavi, Plant and animal small RNA communications between cells and organisms. Nat. Rev. Mol. Cell Biol. 23, 185–203 (2022).

11. Z. Havelda, C. Hornyik, A. Crescenzi, J. Burgyán, J. Burgyan, In Situ Characterization of Cymbidium Ringspot Tombusvirus Infection-Induced Posttranscriptional Gene Silencing in Nicotiana benthamiana. J Virol. 77, 6082–6. (2003).

12. M. Incarbone, A. Zimmermann, P. Hammann, M. Erhardt, F. Michel, P. Dunoyer, Neutralization of mobile antiviral small RNA through peroxisomal import. Nat. Plants. 3, 17094 (2017).

13. T. Xu, L. Zhang, J. Zhen, Y. Fan, C. Zhang, L. Wang, Expressional and regulatory characterization of Arabidopsis RNA-dependent RNA polymerase 1. Planta. 237, 1561–1569 (2013).

14. M. Cao, P. Du, X. Wang, Y.-Q. Yu, Y.-H. Qiu, W. Li, A. Gal-On, C. Zhou, Y. Li, S.-W. Ding, Virus infection triggers widespread silencing of host genes by a distinct class of endogenous siRNAs in Arabidopsis. Proc. Natl. Acad. Sci. 111, 14613–14618 (2014).

15. E. A. Devers, C. A. Brosnan, A. Sarazin, D. Albertini, A. C. Amsler, F. Brioudes, P. E. Jullien, P. Lim, G. Schott, O. Voinnet, Movement and differential consumption of short interfering RNA duplexes underlie mobile RNA interference. Nat. Plants. 6, 789–799 (2020).

16. H. Garcia-Ruiz, A. Takeda, E. J. Chapman, C. M. Sullivan, N. Fahlgren, K. J. Brempelis, J. C. Carrington, Arabidopsis RNA-Dependent RNA Polymerases and Dicer-Like Proteins in Antiviral Defense and Small Interfering RNA Biogenesis during Turnip Mosaic Virus Infection. Plant Cell. 22, 481–496 (2010).

17. H. Garcia-Ruiz, A. Carbonell, J. S. Hoyer, N. Fahlgren, K. B. Gilbert, A. Takeda, A. Giampetruzzi, M. T. Garcia Ruiz, M. G. McGinn, N. Lowery, M. T. Martinez Baladejo, J. C. Carrington, Roles and Programming of Arabidopsis ARGONAUTE Proteins during Turnip Mosaic Virus Infection. PLoS Pathog. 11, 1–27 (2015).

18. S. Li, Z. Zhang, C. Zhou, S. Li, RNA-dependent RNA polymerase 1 delays the accumulation of viroids in infected plants. Mol. Plant Pathol. 22, 1195–1208 (2021).

19. L. Qin, N. Mo, Y. Zhang, T. Muhammad, G. Zhao, Y. Zhang, Y. Liang, CaRDR1, an RNA-Dependent RNA Polymerase Plays a Positive Role in Pepper Resistance against TMV. Front. Plant Sci. 8, 1–13 (2017).

20. L. J. R. Hunter, S. F. Brockington, A. M. Murphy, A. E. Pate, K. Gruden, S. A. MacFarlane, P. Palukaitis, J. P. Carr, RNA-dependent RNA polymerase 1 in potato (Solanum tuberosum) and its relationship to other plant RNA-dependent RNA polymerases. Sci. Rep. 6, 23082 (2016).

21. C. T. Madsen, J. Stephens, C. Hornyik, J. Shaw, D. B. Collinge, C. Lacomme, M. Albrechtsen, Identification and characterization of barley RNA-directed RNA polymerases. Biochim. Biophys. Acta - Gene Regul. Mech. 1789, 375–385 (2009).

22. Y. Peng, J. Yang, X. Li, Y. Zhang, Salicylic Acid: Biosynthesis and Signaling. Annu. Rev. Plant Biol. 72, 761–791 (2021).

23. A. M. Murphy, T. Zhou, J. P. Carr, An update on salicylic acid biosynthesis, its induction and potential exploitation by plant viruses. Curr. Opin. Virol. 42, 8–17 (2020).

24. T. P. Delaney, S. Uknes, B. Vernooij, L. Friedrich, K. Weymann, D. Negrotto, T. Gaffney, M. Gut-Rella, H. Kessmann, E. Ward, J. Ryals, A central role of salicylic acid in plant disease resistance. Science 266, 1247–1250 (1994).

25. M. van Damme, R. P. Huibers, J. Elberse, G. Van den Ackerveken, Arabidopsis DMR6 encodes a putative 2OG-Fe(II) oxygenase that is defense-associated but required for susceptibility to downy mildew. Plant J. 54, 785–793 (2008).

26. J. Chen, W. X. Li, D. Xie, J. R. Peng, S. W. Ding, Viral Virulence Protein Suppresses RNA Silencing–Mediated Defense but Upregulates the Role of MicroRNA in Host Gene Expression. Plant Cell. 16, 1302–1313 (2004).

27. F. Qu, T. Ren, T. J. Morris, The Coat Protein of Turnip Crinkle Virus Suppresses Posttranscriptional Gene Silencing at an Early Initiation Step. J. Virol. 77, 511–522 (2003).

## REFERENCES

1. A. Lampropoulos, Z. Sutikovic, C. Wenzl, I. Maegele, J. U. Lohmann, J. Forner, GreenGate - A Novel, Versatile, and Efficient Cloning System for Plant Transgenesis. PLoS One. 8, e83043 (2013).

2. J. G. Thomson, M. Cook, M. Guttman, J. Smith, R. Thilmony, Novel sul I binary vectors enable an inexpensive foliar selection method in Arabidopsis. BMC Res. Notes. 4, 44 (2011).

3. J. Curaba, X. Chen, Biochemical Activities of Arabidopsis RNA-dependent RNA Polymerase 6. J. Biol. Chem. 283, 3059–3066 (2008).

4. A. Devert, N. Fabre, M. Floris, B. Canard, C. Robaglia, P. Crété, Primer-Dependent and Primer-Independent Initiation of Double Stranded RNA Synthesis by Purified Arabidopsis RNA-Dependent RNA Polymerases RDR2 and RDR6. PLoS One. 10, e0120100 (2015).

5. J. Jiang, C. Patarroyo, D. Garcia Cabanillas, H. Zheng, J.-F. Laliberté, The Vesicle-Forming 6K 2 Protein of Turnip Mosaic Virus Interacts with the COPII Coatomer Sec24a for Viral Systemic Infection. J. Virol. 89, 6695–6710 (2015).

6. M. Laufer, H. Mohammad, E. Maiss, K. Richert-Pöggeler, M. Dall’Ara, C. Ratti, D. Gilmer, S. Liebe, M. Varrelmann, Biological properties of Beet soil-borne mosaic virus and Beet necrotic yellow vein virus cDNA clones produced by isothermal in vitro recombination: Insights for reassortant appearance. Virology. 518, 25–33 (2018).

7. Z. Xie, L. K. Johansen, A. M. Gustafson, K. D. Kasschau, A. D. Lellis, D. Zilberman, S. E. Jacobsen, J. C. Carrington, Genetic and Functional Diversification of Small RNA Pathways in Plants. PLoS Biol. 2, 642–652 (2004).

8. H. Garcia-Ruiz, A. Takeda, E. J. Chapman, C. M. Sullivan, N. Fahlgren, K. J. Brempelis, J. C. Carrington, Arabidopsis RNA-Dependent RNA Polymerases and Dicer-Like Proteins in Antiviral Defense and Small Interfering RNA Biogenesis during Turnip Mosaic Virus Infection. Plant Cell. 22, 481–496 (2010).

9. A. Deleris, J. Gallego-Bartolome, J. Bao, K. D. Kasschau, J. C. Carrington, O. Voinnet, Hierarchical action and inhibition of plant Dicer-like proteins in antiviral defense. Science 313, 68–71 (2006).

10. T. P. Delaney, S. Uknes, B. Vernooij, L. Friedrich, K. Weymann, D. Negrotto, T. Gaffney, M. Gut-Rella, H. Kessmann, E. Ward, J. Ryals, A central role of salicylic acid in plant disease resistance. Science 266, 1247–1250 (1994).

11. M. van Damme, R. P. Huibers, J. Elberse, G. Van den Ackerveken, Arabidopsis DMR6 encodes a putative 2OG-Fe(II) oxygenase that is defense-associated but required for susceptibility to downy mildew. Plant J. 54, 785–793 (2008).

12. M. Incarbone, M. Clavel, B. Monsion, L. Kuhn, H. Scheer, É. Vantard, V. Poignavent, P. Dunoyer, P. Genschik, C. Ritzenthaler, Immunocapture of dsRNA-bound proteins provides insight into Tobacco rattle virus replication complexes and reveals Arabidopsis DRB2 to be a wide-spectrum antiviral effector. Plant Cell. 33, 3402–3420 (2021).

13. T. Pasternak, O. Tietz, K. Rapp, M. Begheldo, R. Nitschke, B. Ruperti, K. Palme, Protocol: an improved and universal procedure for whole-mount immunolocalization in plants. Plant Methods. 11, 50 (2015).

14. J. Schindelin, I. Arganda-Carreras, E. Frise, V. Kaynig, M. Longair, T. Pietzsch, S. Preibisch, C. Rueden, S. Saalfeld, B. Schmid, J.-Y. Tinevez, D. J. White, V. Hartenstein, K. Eliceiri, P. Tomancak, A. Cardona, Fiji: an open-source platform for biological-image analysis. Nat. Methods. 9, 676–682 (2012).

15. Y. Matsushita, T. Usugi, S. Tsuda, Distribution of tomato chlorotic dwarf viroid in floral organs of tomato. Eur. J. Plant Pathol. 130, 441–447 (2011).

16. M. D. Nodine, R. Yadegari, F. E. Tax, RPK1 and TOAD2 Are Two Receptor-like Kinases Redundantly Required for Arabidopsis Embryonic Pattern Formation. Dev. Cell. 12, 943–956 (2007).

17. W. Rozhon, E. Petutschnig, M. Wrzaczek, C. Jonak, Quantification of free and total salicylic acid in plants by solid-phase extraction and isocratic high-performance anion-exchange chromatography. Anal. Bioanal. Chem. 382, 1620–1627 (2005).

18. E. K. Petutschnig, M. Stolze, U. Lipka, M. Kopischke, J. Horlacher, O. Valerius, W. Rozhon, A. A. Gust, B. Kemmerling, B. Poppenberger, G. H. Braus, T. Nürnberger, V. Lipka, A novel Arabidopsis CHITIN ELICITOR RECEPTOR KINASE 1 (CERK1) mutant with enhanced pathogen-induced cell death and altered receptor processing. New Phytol. 204, 955–967 (2014).

19. M. Incarbone, C. Ritzenthaler, P. Dunoyer, Peroxisomal Targeting as a Sensitive Tool to Detect Protein-Small RNA Interactions through in Vivo Piggybacking. Front. Plant Sci. 9, 135 (2018).

20. B. Langmead, S. L. Salzberg, Fast gapped-read alignment with Bowtie 2. Nat. Methods. 9, 357– 359 (2012).

21. F. Ramírez, D. P. Ryan, B. Grüning, V. Bhardwaj, F. Kilpert, A. S. Richter, S. Heyne, F. Dündar, T. Manke, deepTools2: a next generation web server for deep-sequencing data analysis. Nucleic Acids Res. 44, W160–W165 (2016).

22. Y. Liao, G. K. Smyth, W. Shi, featureCounts: an efficient general purpose program for assigning sequence reads to genomic features. Bioinformatics. 30, 923–930 (2014).

23. M. I. Love, W. Huber, S. Anders, Moderated estimation of fold change and dispersion for RNA-seq data with DESeq2. Genome Biol. 15, 550 (2014).

24. H. Wickham, M. Averick, J. Bryan, W. Chang, L. McGowan, R. François, G. Grolemund, A. Hayes, L. Henry, J. Hester, M. Kuhn, T. Pedersen, E. Miller, S. Bache, K. Müller, J. Ooms, D. Robinson, D. Seidel, V. Spinu, K. Takahashi, D. Vaughan, C. Wilke, K. Woo, H. Yutani, Welcome to the Tidyverse. J. Open Source Softw. 4, 1686 (2019).

25. H. Wickham, ggplot2: Elegant Graphics for Data Analysis (Springer-Verlag New York, 2016; https://ggplot2.tidyverse.org).

