## Supplementary material for "A hormone-activated mobile RNAi pathway defends plant stem cells from virus infection": Figures S1 to S10 + legends

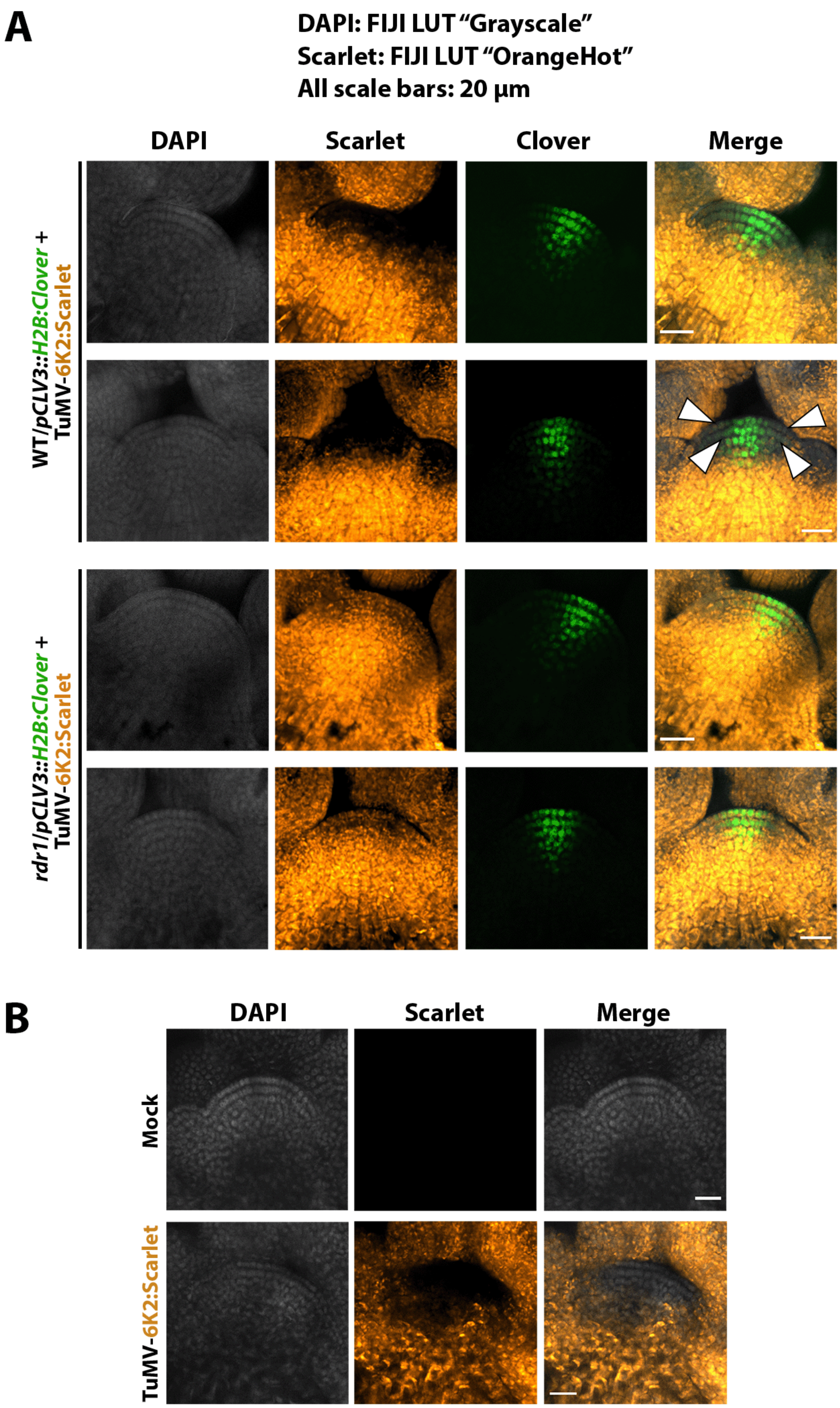


**Figure S1: (A)** Examples of main inflorescence shoot apical meristems (SAMs) from TuMV-6K2:Scarlet-infected WT and *rdr1* mutant transgenic lines expressing nuclear fluorescent reporter H2B:Clover in the stem cell domain of the SAM, through the *pCLV3* promoter, at 16-17 dpi. This color scheme is used for laser confocal microscopy images throughout this study: DAPI fluorescence is in grayscale, Clover is in green (except in Fig. 2D, E), Scarlet fluorescence is in orange-to-yellow (FIJI LUT "OrangeHot", with increasing signal intensity going from orange to yellow). Scale bars always indicate 20 µm. The white arrowheads delineate the "core" virus exclusion zone in WT plants, the two top-most cell layers: L1 and L2. **(B)** As in (A), but WT plants after mock-inoculation (top) or infection with TuMV-6K2:Scarlet.

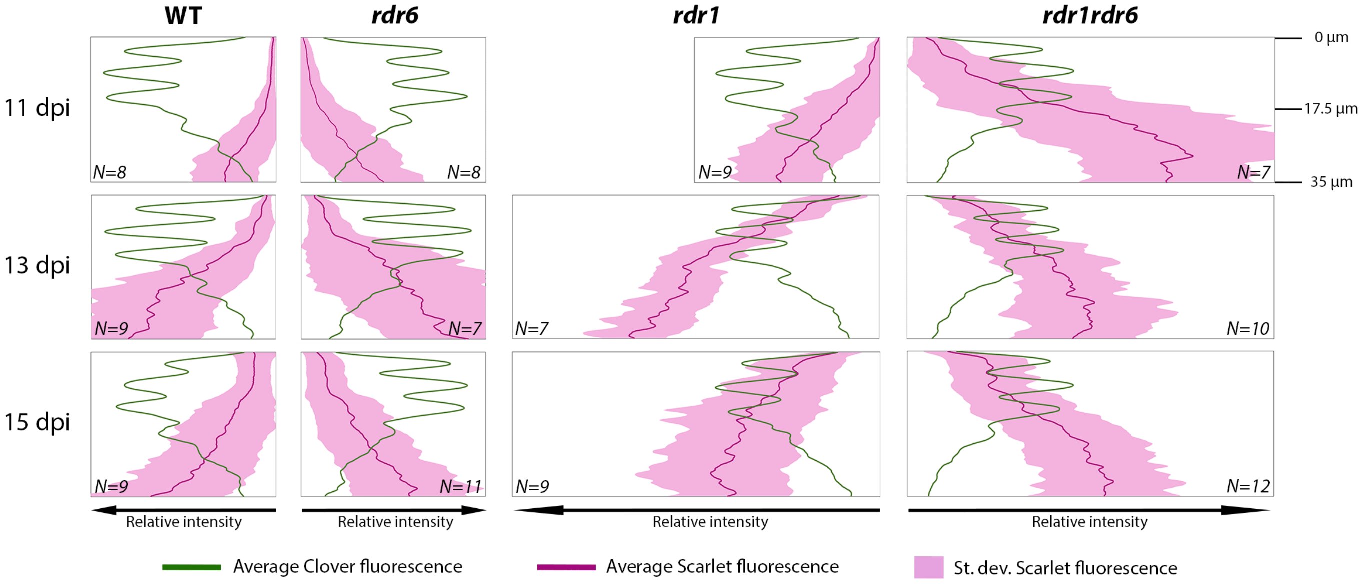


**Figure S2:** Plots showing fluorescence values over time, every 2 days from 11 to 15 days post-inoculation (dpi), in the top 35 µm of the SAM of (from left to right) WT, *rdr6*, *rdr1* and *rdr1rdr6* knock-out mutants (right). The green line indicates the average Clover fluorescence and identifies the L1-L2-L3 cell layers, the dark magenta line indicates the average Scarlet fluorescence (normalized in each sample to the respective Clover values), while the light magenta area indicates the standard deviation of Scarlet fluorescence among the samples. *N* indicates the number of shoot apical meristems analyzed.

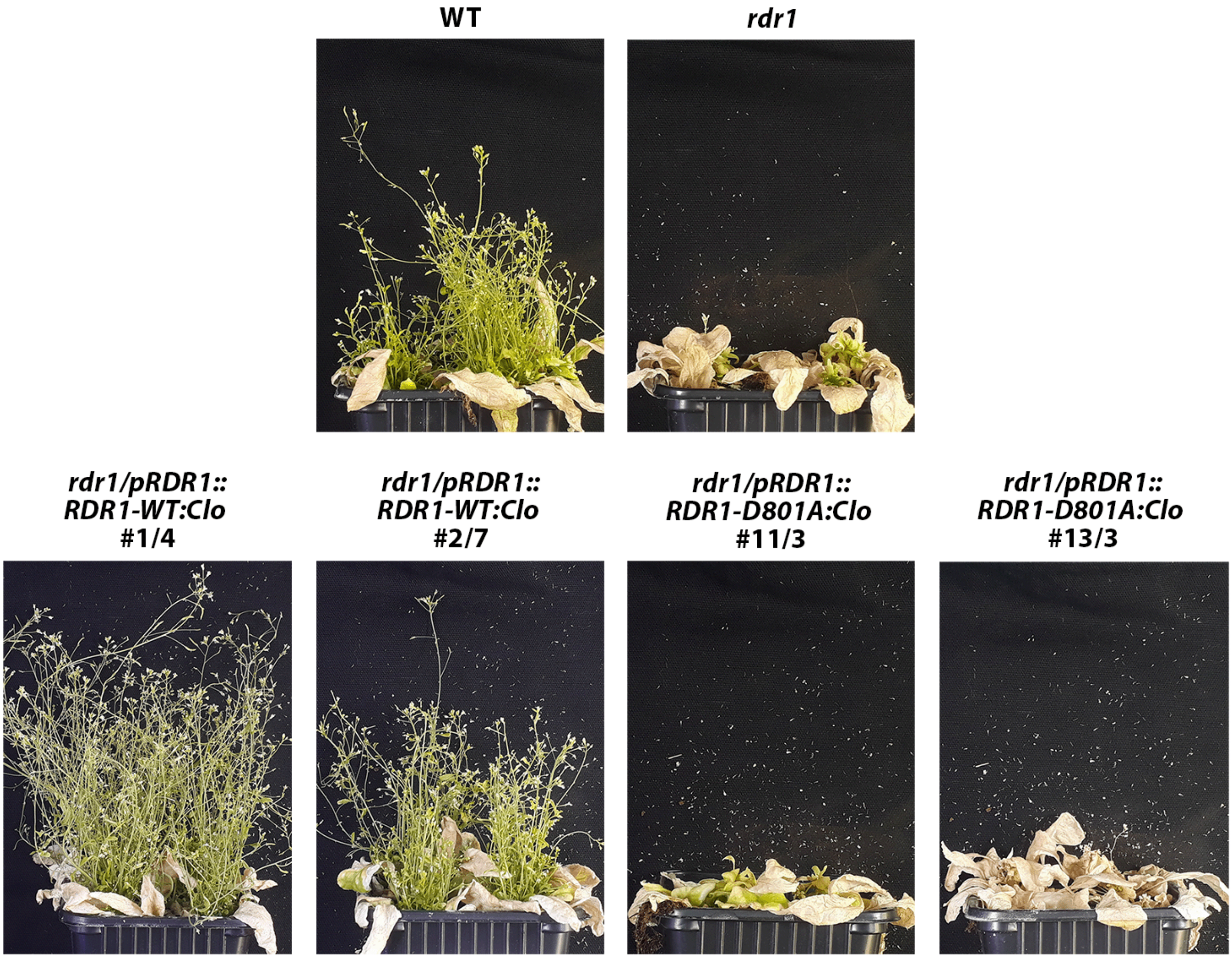


**Figure S3:** Photos of plants one month after inoculation with TuMV-6K2:Scarlet. Top: all infected plants lose apical dominance, but WT generate fertile shoots from axillary meristems, while *rdr1* mutant plants do not. Bottom: *rdr1* mutants complemented with WT (left) or catalytically inactive D801A (right) alleles of Clover-tagged RDR1. Two independent transgenic lines per construct are shown.

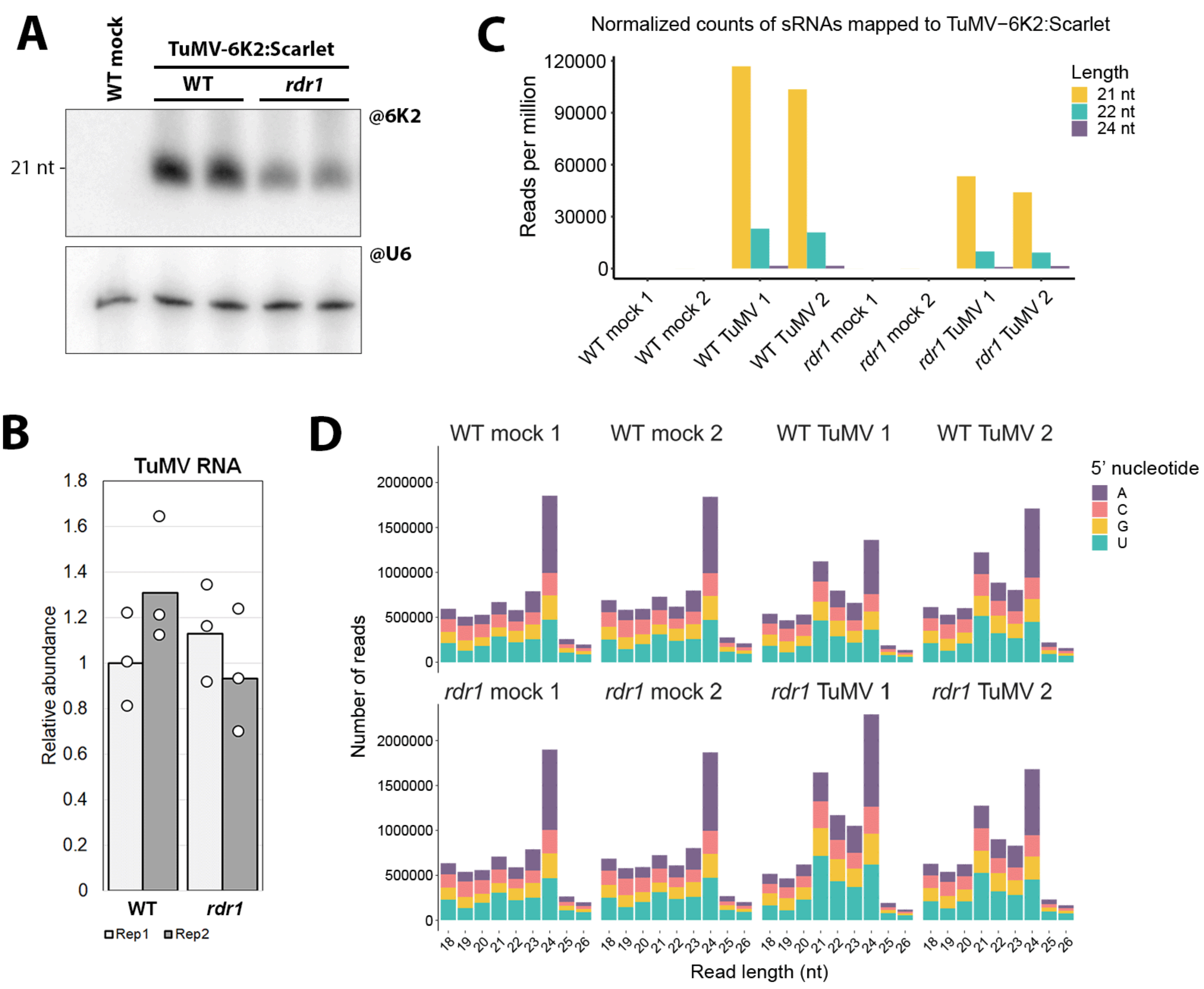


**Figure S4: (A)** PAGE northern blot detection of TuMV 6K2-derived sRNA in systemically infected WT and *rdr1* plants, 11 dpi. Biological duplicates are shown, each sample being a pool of tissue from 4-5 plants. Mock-treated control (plants inoculated with buffer only) is on the left, snoRNA U6 is used as loading control. **(B)** RT-qPCR analysis to quantify TuMV gRNA in the samples described in (A). Each column is a biological replicate, each dot a technical replicate. **(C)** Bar plots showing the number of reads-per-million (divided into 21, 22 and 24 nt in length) of sRNA mapping to TuMV-6K2:Scarlet, as assessed by sRNA sequencing on duplicates of mock- and TuMV-infected apices (meristem and small flower buds) of WT and *rdr1* plants. **(D)** Size distribution (18 to 26 nt length) and 5' nucleotide of the total reads in the experiment described in (C).

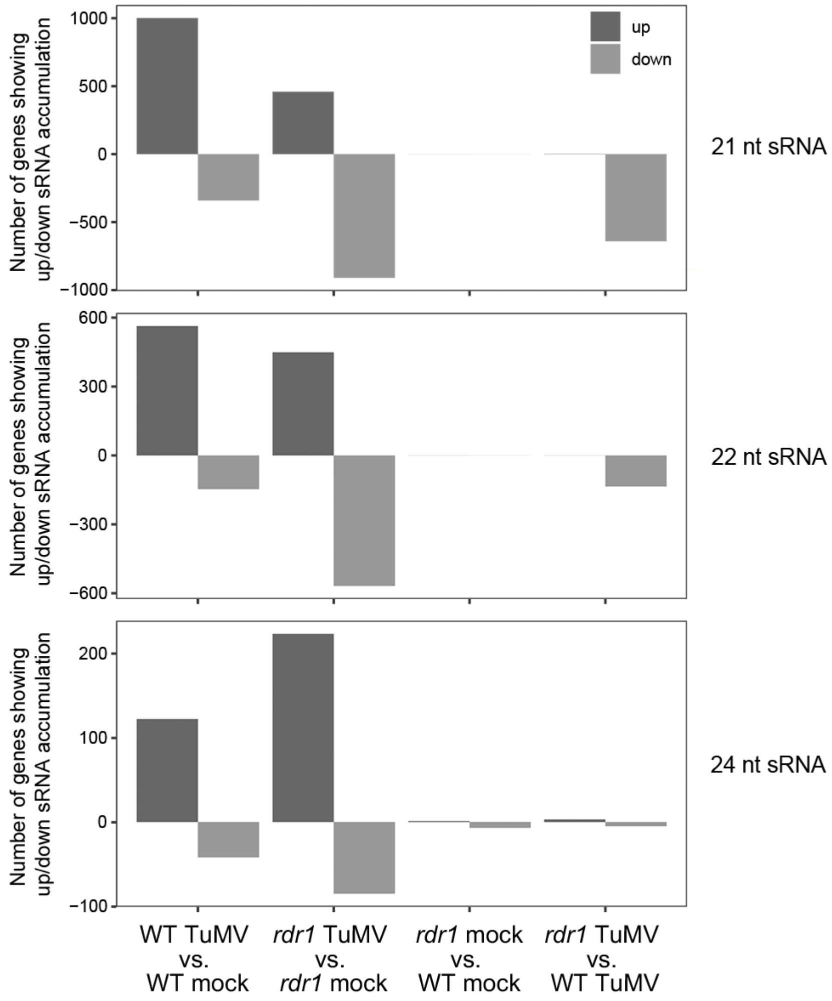


**Figure S5:** Plots showing two-fold increased (up) or decreased (down) accumulation of sRNA mapping to the Arabidopsis genome in mock- and TuMV-infected flower apices of WT and *rdr1* plants, in pairwise comparisons indicated on the bottom horizontal axis. On the vertical axis the plots display the number of Arabidopsis genes from which the up- or down-regulated sRNA are derived. The plots have been divided according to sRNA size: 21nt (top), 22nt (middle) and 24nt (bottom).

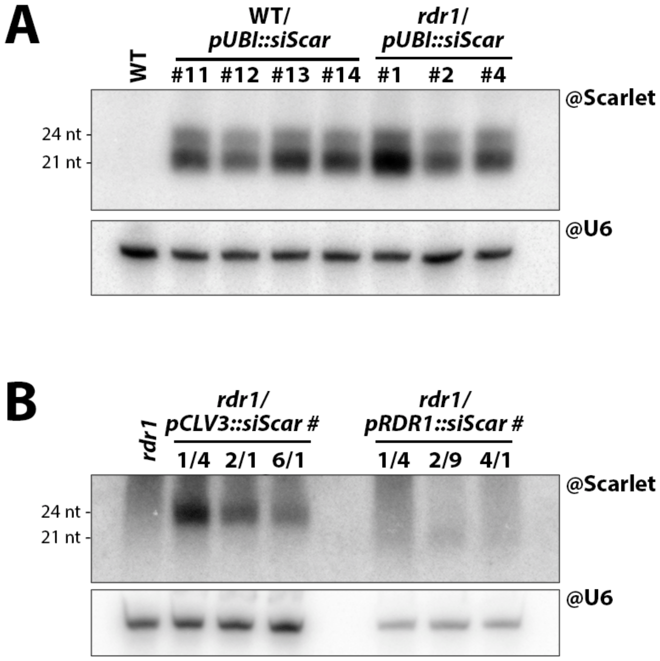


**Figure S6: (A)** PAGE northern blot detection of Scarlet-derived sRNA in seedlings of WT and *rdr1* transgenic lines expressing the *siScar* hairpin construct under the control of the *pUBI* promoter. Each sample is an independent transgenic line. snoRNA U6 is used as loading control. **(B)** As in (A), but in flower apices (meristem and small flower buds) of *rdr1* lines expressing *siScar* through the *pCLV3* promoter (left) and seedlings expressing it through the *pRDR1* promoter. Of note, the *pCLV3*-driven hairpin is processed only into 24 nt-long siRNA.

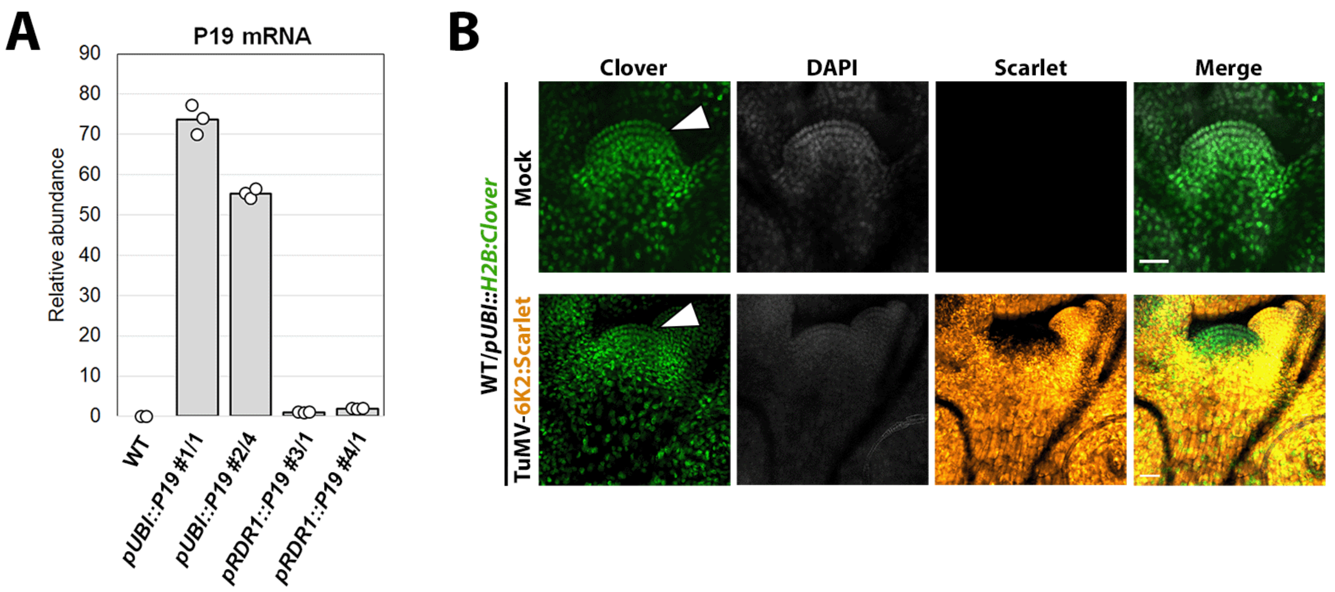


**Figure S7: (A)** RT-qPCR quantification of *P19* expression in upper stems and inflorences of the transgenic lines marked at the bottom, relative to *pRDR1:P19* #3/1. Dots indicate technical replicates. **(B)** Laser confocal microscopy images of meristems from mock and TuMV-6K2:Scarlet-infected plants expressing *H2B:Clover* through the *pUBI* promoter. Expression is ubiquitous, including the L1 and L2 stem cell layers (white arrowheads). DAPI fluorescence in grayscale, Scarlet in orange-to-yellow. Scale bar: 20 µm.

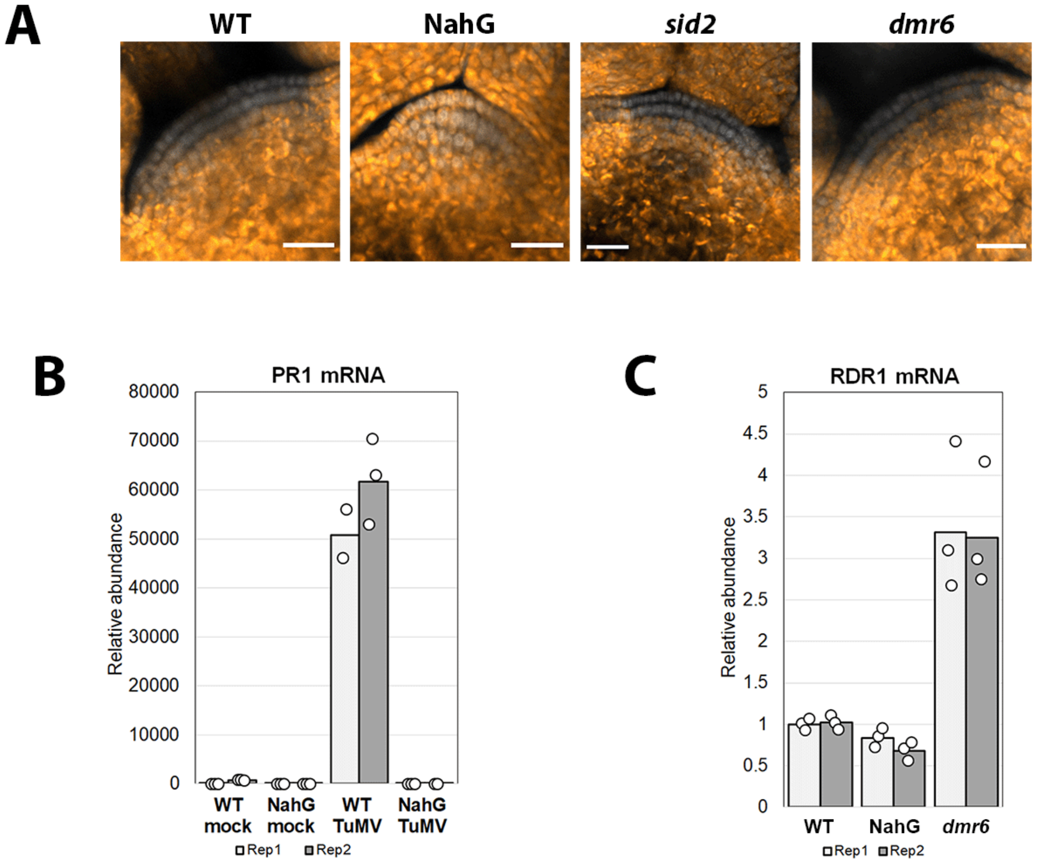


**Figure S8: (A)** Laser confocal microscopy images of meristems from WT, NahG, *sid2* and *dmr6* plants infected with TuMV-6K2:Scarlet. DAPI fluorescence in grayscale, Scarlet in orange-to-yellow. Scale bar: 20 µm. **(B)** RT-qPCR quantification of *PR1* expression in WT and NahG plants after mock or TuMV-6K2:Scarlet inoculation. Each bar is a biological replicate of tissues from 4-5 plants, each dot is a technical replicate. **(C)** RT-qPCR quantification of *RDR1* expression in seedlings of WT, NahG and *dmr6*. Each bar is a biological replicate of 10+ seedlings, each dot is a technical replicate.

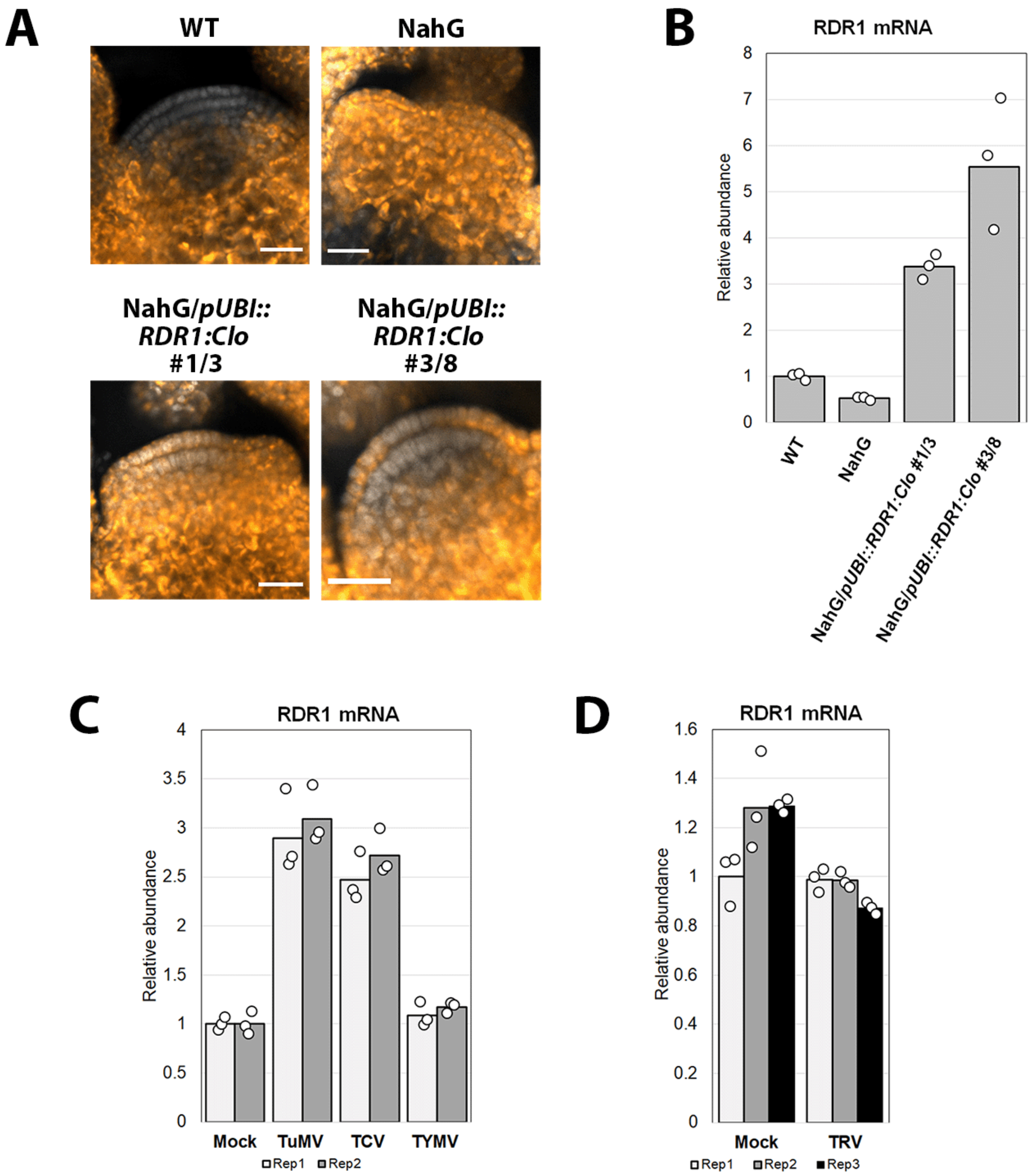


**Figure S9: (A)** Laser confocal microscopy images of meristems from WT, NahG, and two NahG transgenic lines expressing Clover-tagged RDR1 through the *pUBI* promoter, infected with TuMV-6K2:Scarlet. DAPI fluorescence in grayscale, Scarlet in orange-to-yellow. Scale bar: 20 µm. **(B)** RT-qPCR quantification of *RDR1* expression in the plants described in (A). Each dot is a technical replicate. **(C)** RT-qPCR quantification of *RDR1* expression in WT plants systemically infected by TuMV, TCV and TYMV. Each bar is a biological replicate of tissues from 4-5 plants, each dot is a technical replicate. **(D)** As in (C), on plants infected by TRV-Scarlet.

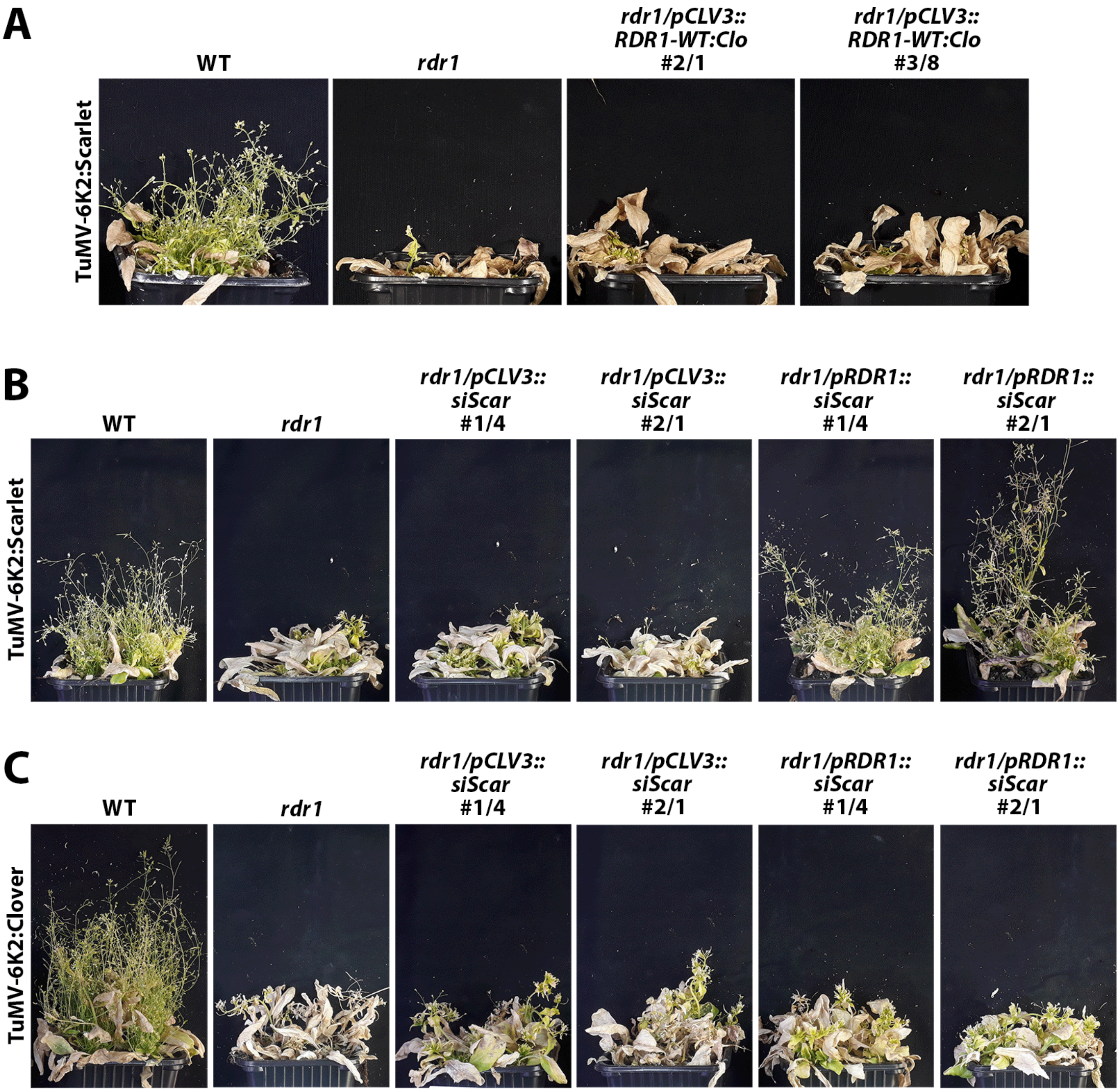


**Figure S10:** **(A)** Photos of plants one month after inoculation with TuMV-6K2:Scarlet. The two *rdr1* transgenic lines on the right express Clover-tagged RDR1 through the *pCLV3* promoter. WT and *rdr1* controls are on the left. **(B)** As in (A), with two *rdr1* transgenic lines expressing the *siScar* construct under the *pCLV3* promoter (middle) or the *pRDR1* promoter (right). **(C)** The lines described in (B), but infected with TuMV-6K2:Clover.
